# Macrophages regulate meiotic initiation and germ cell clearance in the developing ovary

**DOI:** 10.64898/2026.02.28.708733

**Authors:** Xiaowei Gu, Satoko Matsuyama, Shu-Yun Li, Tony DeFalco

## Abstract

Tissue-resident macrophages are increasingly recognized for their roles in promoting organogenesis, yet how macrophages are involved in fetal ovarian development remains unclear. In particular, little is known about ovarian macrophage ontogeny and how it relates to germ cell entry into meiosis and establishment of the oocyte reserve. Here we combine temporally-controlled lineage tracing of yolk-sac erythro-myeloid progenitors, fetal HSC-derived progenitors, and postnatal monocytes to map multi-wave seeding and remodeling of ovarian macrophages across fetal and early postnatal life. We identify three major resident subsets defined by MHCII and CSF1R that display distinct expansion kinetics and persistence, and we show that CCR2-dependent monocyte recruitment is required for efficient maturation of postnatal macrophage populations. Functionally, transient or sustained depletion of CSF1R⁺ fetal macrophages perturbs ovarian vascular growth and triggers precocious meiotic initiation without overt loss of germ cells, leading to persistent, premature meiotic progression. Extending macrophage depletion into late gestation disrupts perinatal physiological germ cell attrition despite rapid postnatal macrophage repopulation. Together, our findings establish ovarian macrophages as stage-specific regulators that couple immune ontogeny to ovarian morphogenesis and germ cell quality control during establishment of the oocyte reserve.

**One Sentence Summary:** Ovarian macrophages are required for the proper timing of germ cell meiotic entry and progression, vascular growth, and for the physiological clearance of germ cells during establishment of the oocyte reserve in perinatal stages.

## Introduction

Organ-resident macrophages have emerged as critical regulators of tissue development, homeostasis, and remodeling, exerting functions beyond immune surveillance to include angiogenesis, stromal patterning, and cellular differentiation (1–3). In reproductive tissues, particularly the ovary, macrophages are abundant, display phenotypic diversity, and undergo dynamic changes across the lifespan (4, 5). However, the ontogeny, proliferative behavior, and functional specialization of ovarian macrophage subsets remain incompletely understood, particularly during fetal and early postnatal windows, when germ cells undergo meiosis and oocyte selection is initiated during establishment of the ovarian reserve.

Tissue-resident macrophages (TRMs) arise through a sequential process involving distinct hematopoietic waves, each characterized by specific timing and progenitor identity (6, 7). In mice, the earliest TRM precursors appear around embryonic day (E) 7.0, originating from primitive erythro-myeloid progenitors (EMPs) in the yolk sac, which primarily generate tissue macrophages without monocyte intermediates. This is followed by a transient hematopoietic phase starting around E8.25, in which a second population of yolk-sac-derived EMPs with broader lineage potential, including monocyte output, migrate to the fetal liver and expand. A third wave initiates around E10.5, when definitive hematopoietic stem cells (HSCs) arise in the aorta-gonad-mesonephros (AGM) region, subsequently colonizing the fetal liver by E12.5. There, they give rise to a range of myeloid and lymphoid cells through steady-state hematopoiesis (8–11), which then migrate to and colonize organs throughout the body. As development proceeds, fetal HSCs seed the emerging bone marrow, forming the foundation for the adult hematopoietic system. Postnatally, circulating monocytes derived from adult bone marrow HSCs can enter tissues and contribute to macrophage pools, particularly in response to injury or remodeling (12, 13). Therefore, this temporal layering of hematopoietic sources establishes a complex ontogeny for TRMs, with each wave leaving lasting imprints on resident macrophage composition and function.

Once established, TRMs acquire specialized phenotypes and functions shaped by local microenvironmental cues, including tissue-derived cytokines, metabolic signals, and stromal interactions. This environmental imprinting confers organ-specific transcriptional and epigenetic identities, enabling macrophages to execute distinct physiological roles (12–14). For instance, yolk-sac-derived microglia sculpt neuronal circuits during development; embryonically seeded Kupffer cells in the liver clear senescent erythrocytes and recycle iron; and yolk-sac-derived Langerhans cells in the epidermis provide frontline barrier immunity (15–17). In the heart and intestine, embryonically-seeded and bone-marrow-derived macrophages coexist and are dynamically replenished, thus coordinating vascular remodeling, tissue repair, and immune homeostasis in response to physiological and inflammatory cues (18, 19). In reproductive organs, fetal HSC-derived testicular macrophages have been shown to regulate Leydig cell steroidogenesis and support spermatogonial niche function (20, 21), while uterine macrophages coordinate implantation, trophoblast invasion, and vascular remodeling during early pregnancy (22, 23). In developing organs, macrophages contribute to tissue morphogenesis, regulate progenitor cell pools, and coordinate vascular and stromal remodeling (24, 25). However, the ovary remains a relatively underexplored site regarding the spatiotemporal organization and tissue-adapted functions of TRMs.

In the fetal ovary, germ cells undergo a tightly timed transition from proliferative oogonia to meiotic oocytes, accompanied by extensive apoptosis and cyst breakdown that shape the oocyte reserve (26). While intrinsic factors like retinoic acid signaling and BMP pathways have been implicated in meiotic entry and oocyte survival (27, 28), the role of immune components, particularly macrophages, in orchestrating these transitions has gained attention. Spatial transcriptomic and single-cell RNA-seq analyses of developing ovaries have revealed that tissue-resident macrophages reside in perivascular niches and reside near germ cells during key developmental windows (29, 30).

Moreover, cytokines such as colony stimulating factor 1 (CSF1) and transforming growth factor beta (TGF-β), known to regulate macrophage survival and activation, are expressed within the ovarian stroma and may signal bidirectionally with germ cells (31–33). Parallel studies in zebrafish models have shown that macrophage activation can influence gonadal sex fate and germ cell survival (34). In mammals, disruptions in macrophage function affect folliculogenesis, angiogenesis, and ovarian aging (35, 36). Additionally, perturbing ovarian TNF-TNFR2 signaling alters oocyte survival and the size of the primordial follicle pool, which points to a conserved but underappreciated immune-germline crosstalk (37). However, direct causal evidence linking macrophage ontogeny to the regulation of meiotic timing and germ cell clearance in the ovary remains limited.

Here, we dissect the developmental origins, expansion kinetics, and functional specialization of ovarian macrophages from embryogenesis through early postnatal stages. Using inducible lineage tracing models targeting yolk-sac-derived (*Csf1r*-creER), fetal HSC-derived (*Kit-*creER), and postnatal monocyte (*Cx3cr1*-creER) lineages, we identify temporally distinct waves of macrophage recruitment and local proliferation. We demonstrate that these macrophages occupy MHCII-defined subsets with divergent proliferative behaviors and tissue persistence. Functional ablation of CSF1R^+^ macrophages resulted in disruptions of germ cell behavior, revealing a requirement for macrophages in preventing premature meiotic entry, promoting physiological germ cell attrition, and coordinating vascular development. Our findings establish ovarian TRMs as multifaceted regulators of early ovarian morphogenesis and oocyte pool homeostasis, expanding the paradigm of developmental immunobiology into the reproductive axis.

## Results

### Ovarian macrophages exhibit temporal control of composition and proliferation

Previous studies have described the presence and phenotypic heterogeneity of ovarian macrophages (38–40), but how their composition and proliferative activity change continuously across fetal and perinatal windows remains poorly understood. To assess the proliferative dynamics of ovarian immune cells across developmental stages, EdU incorporation was evaluated in conjunction with macrophage and immune cell markers. During fetal stages, the ovary possesses two major immune populations: CD45⁺F4/80⁺ macrophages and CD45⁺F4/80⁻ non-macrophage immune cells, which are morphologically smaller and likely represent monocytes (Fig. 1a). Quantitative analyses revealed that at E14.5, CD45⁺F4/80⁺ macrophages constituted the dominant population, far outnumbering CD45⁺F4/80⁻ cells. By E18.5, the number of macrophages remained relatively stable, whereas CD45⁺F4/80⁻ monocyte-like cells underwent a sharp increase in number, reaching levels comparable to those of macrophages (Fig. 1c). These findings suggest a selective proliferation or recruitment of monocytes during late gestation.

**Fig. 1.**
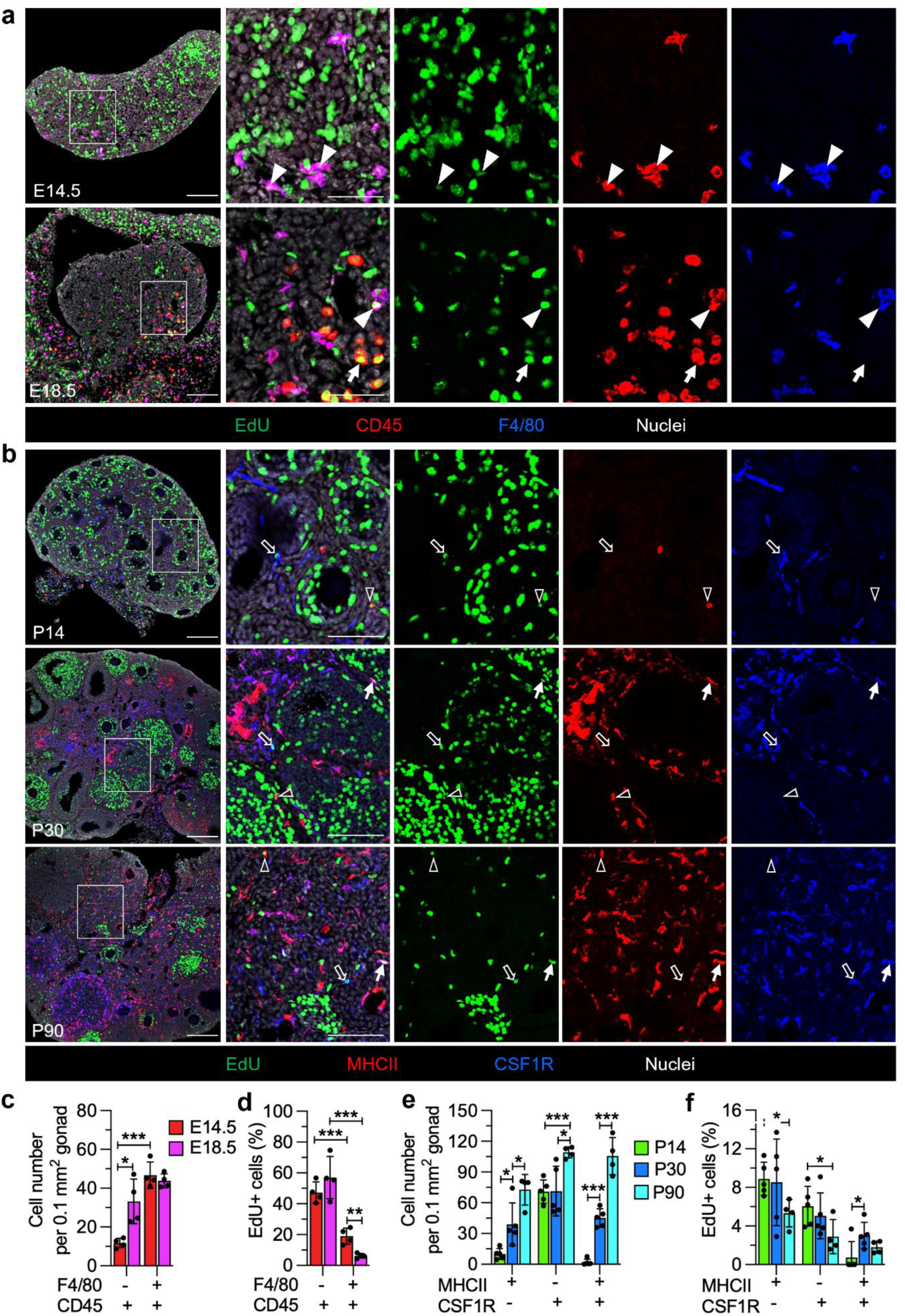
Ovarian macrophage proliferation and subset composition are dynamic during development. (a) Representative sections of E14.5 (*n*=4 independent gonads) and E18.5 (*n*=4) ovaries stained for EdU, CD45, and F4/80. Arrowheads denote EdU⁺CD45⁺F4/80⁺ macrophages; arrows denote EdU⁺CD45⁺F4/80^−^ monocyte-like cells. (b) Representative sections of P14 (*n*=5), P30 (*n*=5), and P90 (*n*=4) ovaries stained for EdU, MHCII, and CSF1R. Black arrowheads indicate EdU⁺MHCII^+^CSF1R^−^ macrophages; black arrows indicate EdU⁺MHCII^−^CSF1R^+^ macrophages; and white arrows indicate EdU⁺MHCII^+^CSF1R^+^ macrophages. (c) Quantification of CD45⁺F4/80⁺ macrophages and CD45⁺F4/80⁻ monocyte-like cells per 0.1 mm² ovarian area at E14.5 and E18.5. (d) Percentage of EdU⁺ cells among macrophages and monocyte-like cells at E14.5 and E18.5. (e) Quantification of MHCII⁻CSF1R⁺, MHCII⁺CSF1R⁺, and MHCII⁺CSF1R⁻ macrophage subsets per 0.1 mm² ovarian area at P14, P30, and P90. (f) Percentage of EdU⁺ cells within each macrophage subset at P14, P30, and P90. Data are shown as mean +/− SD. \**P*<0.05, \*\**P*<0.01, \*\*\**P*<0.001 (two-tailed Student’s *t*-tests). Scale bars, 100 μm (overview) and 50 μm (higher magnification/insets).

In parallel, EdU incorporation data revealed striking differences in proliferative activity between these two populations. At both E14.5 and E18.5, a significantly higher proportion of CD45⁺F4/80⁻ monocyte-like cells were EdU⁺ compared to macrophages, indicating active proliferation. Conversely, the fraction of proliferating macrophages was substantially lower and declined further from E14.5 to E18.5 (Fig. 1d), suggesting that fetal macrophages progressively exit the cell cycle as development advances, while monocytes remain more proliferative.

In the postnatal ovary, macrophages can be divided into three immunophenotypically distinct subpopulations defined by MHCII and CSF1R expression: MHCII⁺CSF1R⁻, MHCII⁺CSF1R⁺, and MHCII⁻CSF1R⁺ (Fig. 1b). At P14, the macrophage pool was dominated by MHCII⁻CSF1R⁺ cells, while both MHCII⁺ subpopulations were present at relatively low levels (Fig. 1e), indicating a predominantly fetal-derived composition at this early postnatal stage. As development progressed, the proportions of MHCII⁺CSF1R⁺ and MHCII⁺CSF1R⁻ macrophages steadily increased at P30 and P90, whereas MHCII⁻CSF1R⁺ cells exhibited only a modest increase by P90, suggesting limited postnatal expansion. EdU incorporation analysis further demonstrated that all three macrophage subtypes were highly proliferative at P14 but showed distinct patterns of age-dependent proliferative decline (Fig. 1f). These data suggest that early postnatal macrophage expansion is driven by active proliferation.

### Csf1r^+^ fetal definitive progenitors give rise to fetal ovarian macrophages

To elucidate the origins and contributions of ovarian macrophages during development, we employed a *Csf1r*-creER; *Rosa*-Tomato lineage tracing system. Pregnant mice were administered a single dose of 4-hydroxytamoxifen (4-OHT) at three critical embryonic time points: E8.5, exclusively targeting yolk sac (YS)-derived erythro-myeloid progenitors (EMPs) (41, 42); E10.5, targeting both YS EMPs and fetal hematopoietic stem cell (HSC)-derived progenitors (21); or E12.5, mainly targeting fetal HSC-derived progenitors (21) (Fig. 2a). This strategy allowed us to evaluate the contribution of labeled progenitors to the macrophage population at E18.5, representing the late fetal ovary, as well as their persistence and distribution in the postnatal and adult ovary at P30 and P90, respectively.

**Fig. 2.**
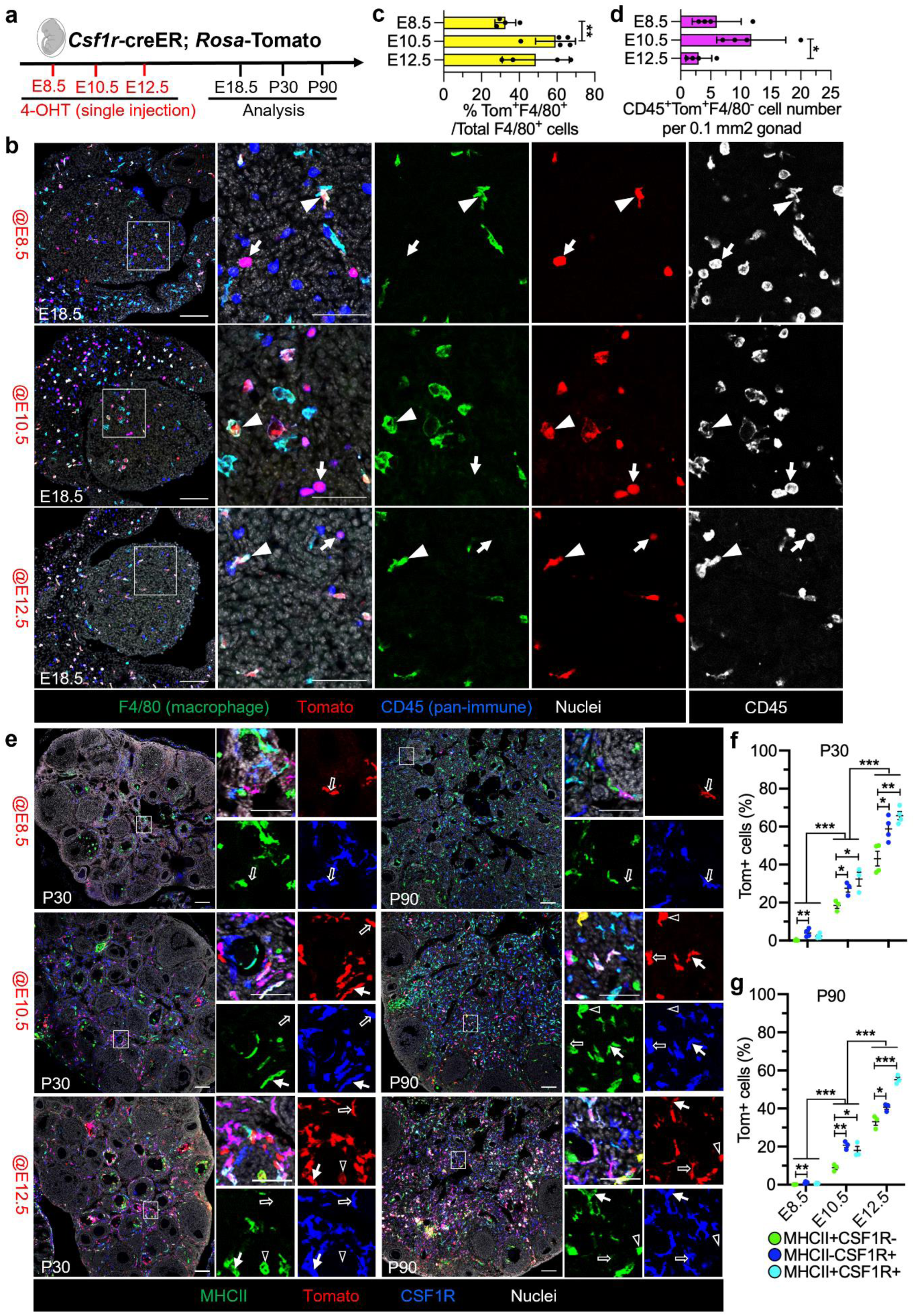
*Csf1r*⁺ fetal definitive progenitors give rise to fetal and postnatal ovarian macrophages. (a) Scheme of the *Csf1r*-creER; *Rosa*-Tomato lineage-tracing strategy. (b) Representative E18.5 ovarian images (*n*=5) showing Tomato-labeled CD45⁺F4/80⁺ macrophages and CD45⁺F4/80⁻ monocyte-like cells in each induction group. Arrowheads, Tomato⁺ macrophages; arrows, Tomato⁺ monocyte-like cells. (c) Percentage of Tomato⁺ cells among total F4/80⁺ macrophages at E18.5 following the indicated induction times. (d) Number of Tomato⁺CD45⁺ F4/80⁻ monocyte-like cells per 0.1 mm² ovarian area at E18.5. (e) Representative P30 and P90 ovaries (*n*=4) stained for MHCII, CSF1R, and Tomato, illustrating the contribution of *Csf1r*^+^ lineage cells to MHCII⁻CSF1R⁺, MHCII⁺CSF1R⁺, and MHCII⁺CSF1R⁻ macrophage subsets after induction at E8.5, E10.5 or E12.5. Black arrows denote Tomato⁺MHCII^−^CSF1R^+^ macrophages; white arrows denote Tomato⁺MHCII^+^CSF1R^+^ macrophages; and black arrowheads denote Tomato⁺MHCII^+^CSF1R^−^macrophages. (f and g) Percentages of Tomato⁺ cells within each macrophage subset at P30 (f) and P90 (g). Data are shown as mean +/− SD. \**P*<0.05, \*\**P*<0.01, \*\*\**P*<0.001 (two-tailed Student’s *t*-tests). Scale bars, 100 μm (overview) and 50 μm (higher magnification/insets).

Immunofluorescence imaging of E18.5 ovaries revealed the presence of Tomato-labeled macrophages (CD45^+^F4/80^+^Tom^+^) across all induction groups, with abundance differing based on the timing of 4-OHT administration (Fig. 2b). Quantitative analysis indicated that E10.5 induction produced the highest proportion of Tomato^+^ macrophages (∼60%), followed by E12.5 (∼50%), and E8.5 (∼30%) (Fig. 2c). In addition to macrophages, CD45^+^F4/80^−^Tom^+^ monocytes were also detected, with their density in E10.5-labeled ovaries significantly exceeding that of E12.5-labeled ovaries but not differing significantly from E8.5-labeled ovaries (Fig. 2d). These findings highlight the dual contributions of YS-derived progenitors and fetal hematopoietic stem cells (HSCs) to the ovarian macrophage population at late fetal stages.

Postnatal analysis at P30 and P90 showed that embryonically-labeled macrophages persisted into adulthood, with notable differences in abundance based on the timing of labeling (Fig. 2e). At P30, macrophages derived from E12.5-labeled progenitors were the most abundant, contributing 45% of MHCII^+^CSF1R^−^, 60% of MHCII^−^CSF1R^+^, and 65% of MHCII^+^CSF1R^+^ subpopulations (Fig. 2f). In contrast, progenitors labeled at E10.5 contributed between 20% to 35% of ovarian macrophages, while those labeled at E8.5 represented less than 5%. At P90, a similar trend was observed, with progenitors originating from E12.5 accounting for 30-45%, those from E10.5 representing 10-20%, and those from E8.5 approaching negligible levels (Fig. 2g).

### HSC-derived macrophages eventually replace EMP-derived macrophages in the adult ovary

Because *Csf1r*-based inducible fate mapping labels not only fetal HSC-derived progenitors but also differentiated macrophages (43, 44), it is difficult to unambiguously resolve the specific contribution of HSC-derived lineages to ovarian macrophages using this approach alone. To further explore the developmental roles of hematopoietic progenitors with respect to ovarian macrophages, we utilized a *Kit*-creER; *Rosa*-Tomato lineage tracing system, which allowed for accurate tracking of hematopoietic progenitors at various gestational stages (21, 45). Induction at E8.5 marked early YS-derived progenitors, whereas inductions at E10.5 and E12.5 mainly labeled HSCs and fetal liver progenitors that contribute to definitive hematopoiesis (21). Consequently, pregnant mice received a single administration of 4-OHT at E8.5, E10.5, or E12.5 (Fig. 3a). The labeled cells were subsequently analyzed at E18.5 to evaluate their contribution to fetal ovarian macrophages and during postnatal and adult stages (P30, P60, and P90) to investigate their long-term presence and distribution within macrophage subpopulations.

**Fig. 3.**
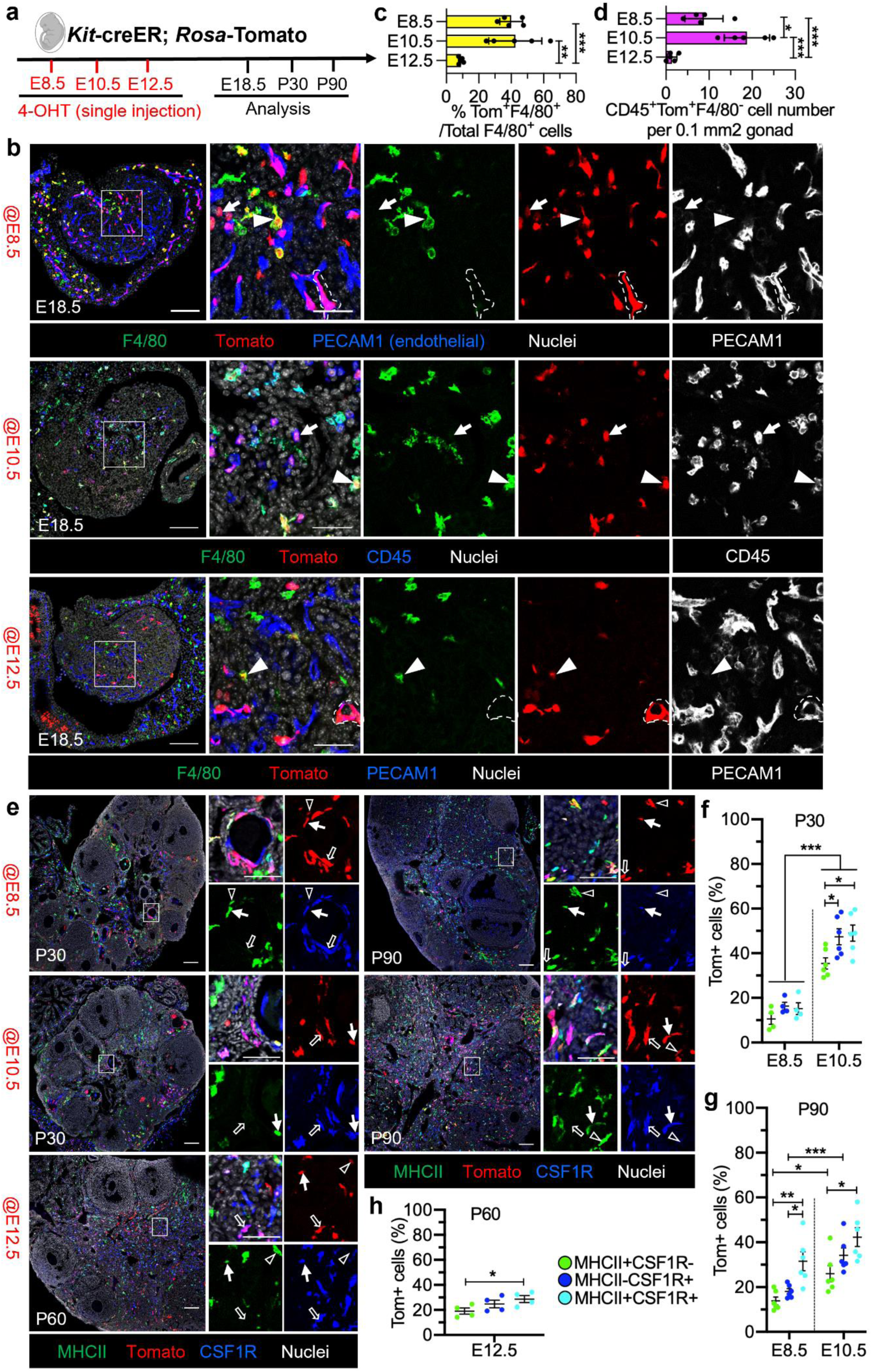
Fetal HSCs progressively replace EMPs to generate adult ovarian macrophages. (a) Scheme of the *Kit*-creER; *Rosa*-Tomato lineage-tracing strategy. (b) Representative E18.5 ovarian images (*n*=5) showing Tomato-labeled CD45⁺F4/80⁺ macrophages, CD45⁺F4/80⁻ monocyte-like cells, and PECAM1⁺ endothelial cells after induction at the indicated stages. Arrowheads denote Tomato⁺ macrophages; arrows denote Tomato⁺ monocyte-like cells; dashed outlines mark Tomato⁺PECAM1⁺ endothelial cells. (c) Percentage of Tomato⁺ cells among total F4/80⁺ macrophages at E18.5 in each induction group. (d) Number of Tomato⁺CD45⁺F4/80⁻ monocyte-like cells per 0.1 mm² ovarian area at E18.5. (e) Representative P30 (*n*=4), P60 (*n*=4), and P90 (*n*=6) ovaries stained for MHCII, CSF1R, and Tomato, illustrating the distribution of *Kit*-lineage cells among MHCII⁻CSF1R⁺, MHCII⁺CSF1R⁺, and MHCII⁺CSF1R⁻ macrophage subsets following induction at E8.5, E10.5 or E12.5. Black arrows denote Tomato⁺MHCII^−^CSF1R^+^ macrophages; white arrows denote Tomato⁺MHCII^+^CSF1R^+^ macrophages; and black arrowheads denote Tomato⁺MHCII^+^CSF1R^−^ macrophages. (f and g) Percentages of Tomato⁺ cells within each macrophage subset at P30 (f) and P90 (g) for E8.5 and E10.5 inductions. (h) Contribution of E12.5-labeled cells to MHCII⁻CSF1R⁺, MHCII⁺CSF1R⁺, and MHCII⁺CSF1R⁻ macrophages at P60. Data are shown as mean +/− SD. \**P*<0.05, \*\**P*<0.01, \*\*\**P*<0.001 (two-tailed Student’s *t*-tests). Scale bars, 100 μm (overview) and 50 μm (higher magnification/insets).

Immunofluorescence imaging of E18.5 ovaries demonstrated distinct contributions of Tomato-labeled cells (CD45^+^F4/80^+^Tom^+^ macrophages, CD45^+^F4/80^−^Tom^+^ monocytes, and PECAM1^+^Tom^+^ endothelial cells) among the induction groups, based on the timing of 4-OHT administration (Fig. 3b). Both E8.5 and E10.5 inductions exhibited comparable macrophage labeling efficiency, which was markedly higher than the minimal macrophage labeling noted for E12.5 induction (Fig. 3c).

Regarding CD45^+^F4/80^−^Tom^+^ monocyte labeling, E10.5 induction yielded the highest density of labeled monocytes, significantly surpassing that of E8.5 induction, while E12.5 labeled monocytes were rarely observed (Fig. 3d). Furthermore, a subset of PECAM1^+^ endothelial cells was identified with E8.5 induction, while E12.5 induction primarily labeled endothelial cells. These results further support the idea that late fetal ovarian macrophages arise from both YS-derived progenitors and fetal HSCs, whereas monocytes are chiefly derived from fetal HSCs, underscoring the distinct origins of these immune cell populations throughout ovarian development.

We discovered that E10.5-labeled macrophages were predominant across all subpopulations at P30 and P90, particularly among MHCII^+^CSF1R^+^ ovarian macrophages. Conversely, E8.5-labeled macrophages represented roughly 20% of the overall macrophage population at these time points, which is significantly less than the contribution of E10.5-labeled macrophages (Fig. 3e-g). While induction at E12.5 had an insignificant impact on ovarian macrophage populations at E18.5, it accounted for about 30% of ovarian macrophages at P60, with a notably greater proportion of labeled MHCII^+^CSF1R^+^ macrophages compared to MHCII^−^CSF1R^+^ macrophages (Fig. 3e and 3h). These results suggest that E10.5-derived fetal HSCs are the primary source of ovarian macrophage populations in adulthood, whereas E12.5-labeled progenitors may migrate into the bone marrow, acting as bone-marrow-derived progenitors and providing a delayed but meaningful contribution to ovarian macrophages.

### Postnatal blood monocytes progressively differentiate into ovarian macrophages

Given that *Cx3cr1*-creER is widely used to trace monocytes and tissue-resident macrophages (46), we employed the *Cx3cr1*-creER; *Rosa*-Tomato lineage tracing model to investigate the contributions of *Cx3cr1*^+^ progenitors to ovarian macrophage populations. *Cx3cr1*-expressing cells were labeled via induction with 4-OHT at E12.5 to label fetal populations or at P4 and P5 with tamoxifen (TAM) to label early postnatal populations (Fig. 4a and 4d).

**Fig. 4.**
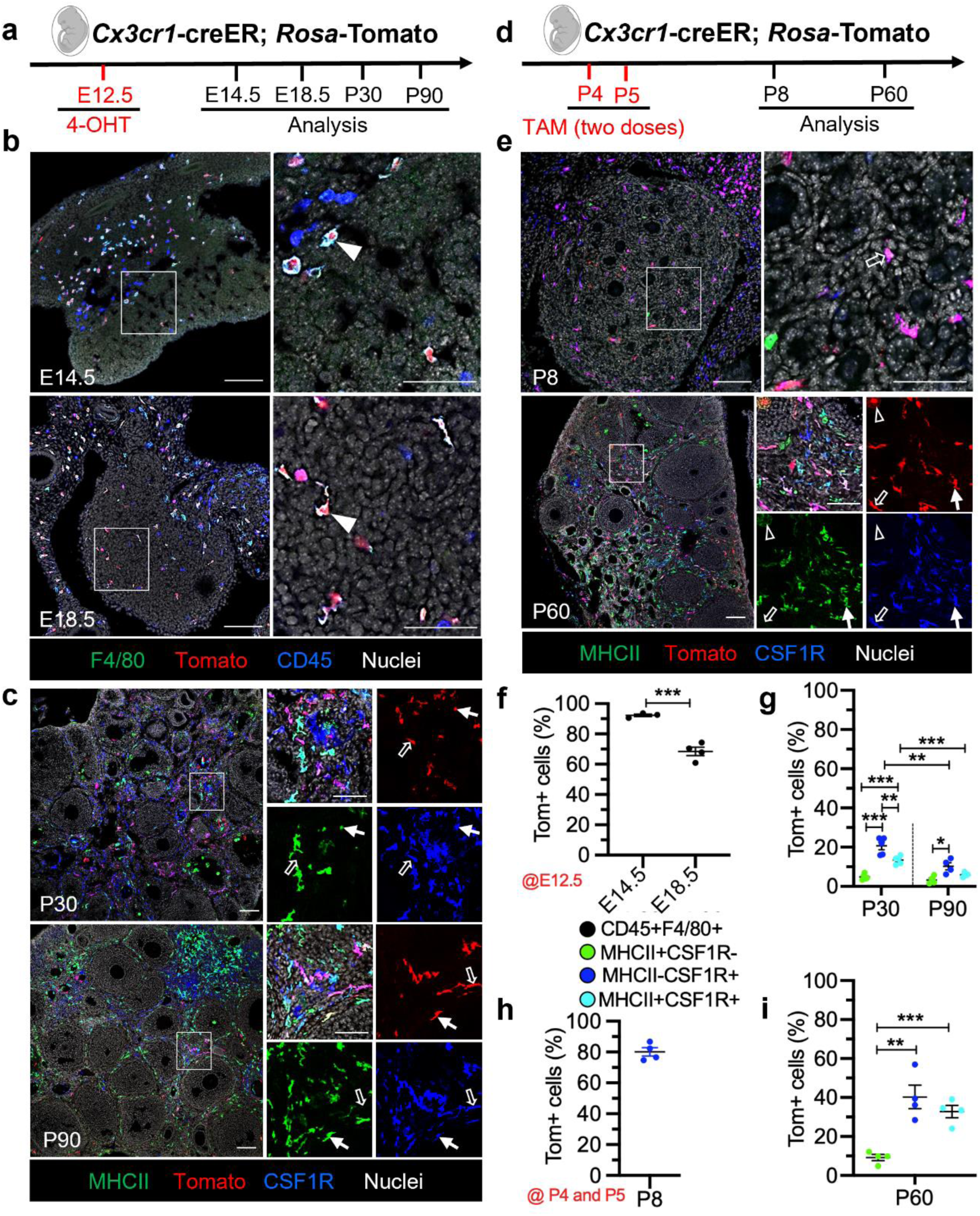
*Cx3cr1*⁺ embryonic and postnatal progenitors contribute to ovarian macrophage subsets. (a) Scheme of embryonic *Cx3cr1*-creER; *Rosa*-Tomato labeling. (b) Representative E14.5 (*n*=3) and E18.5 (*n*=4) ovarian sections stained for F4/80, CD45, and Tomato following E12.5 induction. Arrowheads indicate Tomato⁺CD45^+^F4/80^+^ macrophages. (c) Representative P30 (*n*=5) and P90 (*n*=4) ovarian sections stained for MHCII, CSF1R, and Tomato, showing the persistence of embryonically labeled macrophages within MHCII⁻CSF1R⁺, MHCII⁺CSF1R⁺, and MHCII⁺CSF1R⁻ subsets. Black arrows denote Tomato⁺MHCII^−^CSF1R^+^ macrophages and white arrows denote Tomato⁺MHCII^+^CSF1R^+^ macrophages. (d) Scheme of postnatal *Cx3cr1*-creER*; Rosa*-Tomato labeling. Pups received TAM at P4 and P5 and ovaries were analyzed at P8 and P60. (e) Representative P8 (*n*=4) and P60 (*n*=4) ovarian sections from postnatally induced animals stained for MHCII, CSF1R, and Tomato. Black arrows denote Tomato⁺MHCII^−^CSF1R^+^ macrophages, white arrows denote Tomato⁺MHCII^+^CSF1R^+^ macrophages, and black arrowheads denote Tomato⁺MHCII^+^CSF1R^−^ macrophages. (f) Quantification of percent Tomato⁺ cells among CD45⁺F4/80⁺ macrophages at E14.5 and E18.5 after E12.5 induction. (g) Percentages of Tomato⁺ cells within each macrophage subset at P30 and P90. (h and i) Percentages of Tomato⁺ cells within MHCII⁻CSF1R⁺ macrophages at P8 (h), and within MHCII⁻CSF1R⁺, MHCII⁺CSF1R⁺, and MHCII⁺CSF1R⁻ macrophages at P60 (i) after P4/P5 induction. Data are shown as mean +/− SD. \**P*<0.05, \*\**P*<0.01, \*\*\**P*<0.001 (two-tailed Student’s *t*-tests). Scale bars, 100 μm (overview) and 50 μm (higher magnification/insets).

Upon induction at E12.5, we observed that nearly 90% of CD45^+^F4/80^+^ macrophages at E14.5 were Tomato-labeled, with this percentage diminishing to 70% by E18.5, reflecting a progressive decline in macrophage labeling as development advanced (Fig. 4b and 4f). By P30, fewer than 20% of each macrophage subpopulation was Tomato-labeled, and by P90, this proportion further declined to less than 10% (Fig. 4c and 4g). While the contribution of E12.5-labeled cells to both MHCII^+^CSF1R^+^ and MHCII^−^CSF1R^+^ macrophages was higher than their contribution to MHCII^+^CSF1R^−^ macrophages, all subpopulations exhibited a significant reduction in labeling from P30 to P90 (Fig. 4g). These results indicate that E12.5-labeled ovarian macrophages begin to undergo a significant decrease in number during fetal development, with very few persisting into adulthood across all macrophage subpopulations.

Following TAM administration at P4 and P5, 80% of CSF1R^+^ macrophages in P8 ovaries exhibited Tomato expression, demonstrating effective labeling of early postnatal macrophages at this stage. Importantly, MHCII^+^ macrophages were absent in the ovary at this time, underscoring the early developmental phase of the macrophage population (Fig. 4e and 4h). By P60, a small fraction (∼10%) of MHCII^+^CSF1R^−^ macrophages was labeled, indicating that *Cx3cr1*-creER-labeled postnatal progenitors contributed to MHCII^+^ ovarian macrophages. Meanwhile, ∼30% of MHCII^+^CSF1R^+^ macrophages and ∼40% of MHCII^−^CSF1R^+^ macrophages were Tomato-labeled. However, the labeling efficiency at P60 was markedly lower compared to the high Tomato labeling observed in CSF1R^+^ macrophages at P7 (Fig. 4e and 4i). This may be due to the lower recombination rate of *Cx3cr1*-creER in myeloid cells, particularly those contributing to MHCII^+^ subpopulations. These findings indicate that *Cx3cr1*-creER-labeled early postnatal progenitors contribute to the establishment of ovarian macrophage populations, including MHCII^+^ macrophages, over time.

### Ccr2 is required for the recruitment and differentiation of monocyte-derived macrophages in the postnatal ovary

Given CCR2 is a key chemokine receptor that governs the egress of classical monocytes from the bone marrow and their recruitment into peripheral tissues (47, 48), we utilized the *Ccr2^GFP^* knock- in/knockout model to assess whether CCR2-dependent monocyte trafficking shapes the postnatal ovarian macrophage pool. Therefore, we compared *Ccr2^GFP/+^* (control) and *Ccr2^GFP/GFP^* (i.e., *Ccr2* KO) ovaries at P30 and P60. In control ovaries, GFP⁺ cells primarily contributed to MHCII⁺CSF1R⁻ and MHCII⁺CSF1R⁺ macrophage subpopulations, with the highest contribution observed in the MHCII⁺CSF1R⁻ subset, and minimal contribution to the MHCII⁻CSF1R⁺ population (Fig. 5a-d). This pattern suggests that in the adult ovary, MHCII⁺CSF1R⁻ and MHCII⁺CSF1R⁺ macrophages are largely derived from blood monocytes.

**Fig. 5.**
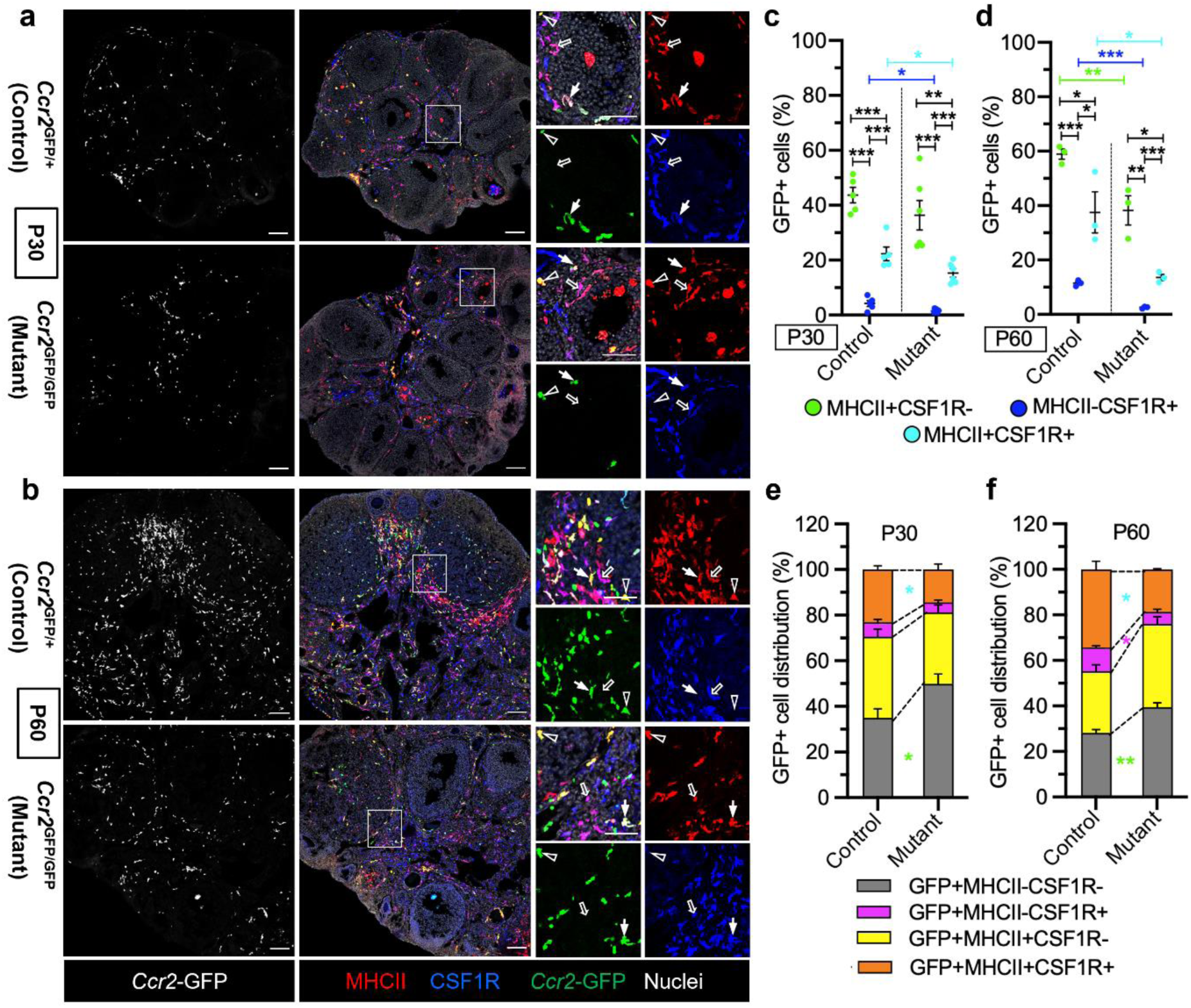
*Ccr2* is required for recruitment and differentiation of monocyte-derived ovarian macrophages. (a) Representative P30 ovarian sections from *Ccr2^GFP/+^*control (*n*=6) and *Ccr2^GFP/GFP^* KO (*n*=6) mice stained for MHCII and CSF1R. Black arrows indicate GFP-negative MHCII^−^CSF1R^+^ macrophages; white arrows indicate GFP-expressing MHCII^+^CSF1R^+^ macrophages; and black arrowheads indicate GFP-expressing MHCII^+^CSF1R^−^ macrophages. (b) Representative P60 ovarian sections from *Ccr2^GFP/+^* control (*n*=4) and *Ccr2^GFP/GFP^* KO (*n*=4) mice stained for MHCII and CSF1R. (c and d) Percentages of GFP⁺ cells within MHCII⁻CSF1R⁺, MHCII⁺CSF1R⁺, and MHCII⁺CSF1R⁻ macrophage subsets in control and KO ovaries at P30 (c) and P60 (d). (e and f) Proportional distribution of GFP⁺ cells among MHCII⁻CSF1R⁻, MHCII⁻CSF1R⁺, MHCII⁺CSF1R⁺, and MHCII⁺CSF1R⁻ populations at P30 (e) and P60 (f). Data are shown as mean +/− SD. \**P*<0.05, \*\**P*<0.01, \*\*\**P*<0.001 (two-tailed Student’s *t*-tests). Scale bars, 100 μm (overview) and 50 μm (higher magnification/insets).

*Ccr2* KO ovaries exhibited a marked reduction in GFP⁺ macrophages exhibiting MHCII⁺CSF1R⁺ and MHCII⁻CSF1R⁺ phenotypes at P30, and a significant reduction in all three macrophage subpopulations by P60 (Fig. 5a-d). In addition, there was a corresponding increase in GFP⁺MHCII⁻CSF1R⁻ cells in *Ccr2* KO ovaries at both P30 and P60, indicative of arrested or incomplete differentiation of CCR2^+^ monocytes (Fig. 5e and 5f). These results demonstrate that CCR2 is essential not only for the recruitment of monocytes but also for their subsequent maturation into fully differentiated ovarian macrophages during postnatal development.

### CSF1R-dependent fetal ovarian macrophages regulate vascular development and meiotic initiation

CSF1R signaling is essential for the early development and migration of yolk-sac-derived macrophage progenitors (49); therefore, we treated pregnant mice with anti-CSF1R antibody (or control IgG antibody) at E6.5 to deplete yolk-sac-derived macrophages in vivo, as previously reported (9, 21, 50). Fetal ovaries were then analyzed at E13.5, E14.5, and E18.5 (Fig. 6a, 6c and 6e). At E13.5, F4/80^+^ macrophages were markedly reduced in anti-CSF1R-treated ovaries, while the number of CD11b^+^CD45^+^ monocytes increased, suggesting that CSF1R blockade impaired the differentiation of monocytes into macrophages (Fig. 6a), and resulted in monocyte accumulation. Immunostaining with PECAM1 revealed a slight reduction in vascular density (Fig. 6a), indicating that macrophages contribute to vascular expansion. Gene expression analysis further confirmed decreased expression of both macrophage-related genes (*Adgre1, Cx3cr1, Csf1r, Mrc1*) and the endothelial marker *Cdh5* in anti-CSF1R-treated ovaries (Fig. 6b), supporting the immunohistological findings.

**Fig. 6.**
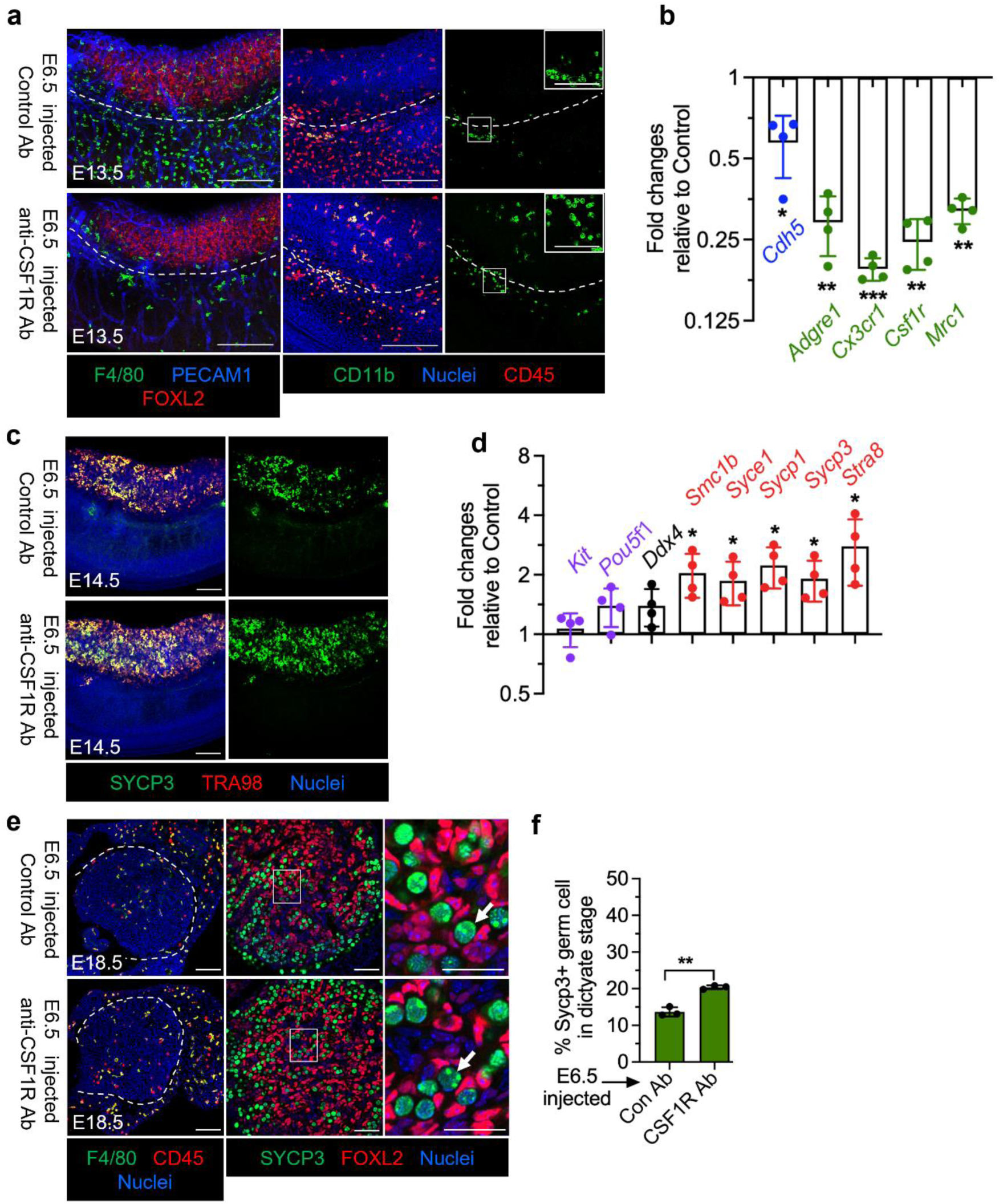
Antibody-mediated depletion of CSF1R⁺ fetal macrophages disrupts vascular growth and induces premature meiotic initiation. (a) Representative whole-mount images of E13.5 ovaries from C57BL/6J embryos exposed at E6.5 to either control rat IgG2a antibody or anti-CSF1R blocking antibody to deplete YS-derived macrophages. For F4/80, PECAM1, and FOXL2 staining, *n*=5 control and *n*=6 anti-CSF1R-treated independent gonads; for CD11b and CD45 staining, *n*=3 control and *n*=3 anti-CSF1R-treated independent gonads. Insets show higher-magnification views of boxed regions; dashed lines mark the gonad-mesonephros boundary. (b) qRT-PCR analysis of macrophage-associated genes (*Adgre1, Cx3cr1, Csf1r, Mrc1*) and endothelial marker *Cdh5*, showing fold change in gene expression in E13.5 anti-CSF1R-treated ovaries versus controls. (c) Representative E14.5 sections stained for SYCP3 and TRA98 showing increased meiotic germ cells in anti-CSF1R-treated ovaries (*n*=4) compared with control ovaries (*n*=4). (d) qRT-PCR analysis of germ cell and meiotic genes (*Kit, Pou5f1, Ddx4, Stra8, Sycp1, Sycp3, Syce1, Smc1b*), showing fold change in gene expression in E14.5 anti-CSF1R-treated ovaries versus controls. (e) Representative E18.5 sections stained for F4/80 and CD45, and for SYCP3 and FOXL2, showing recovery of F4/80⁺ macrophages and advanced meiotic progression in anti-CSF1R-treated ovaries compared with control ovaries. For F4/80 and CD45 staining, *n*=4 control and *n*=4 anti-CSF1R-treated independent gonads; for SYCP3 and FOXL2 staining, *n*=3 control and *n*=3 anti-CSF1R-treated independent gonads. Arrows indicate SYCP3⁺ dictyate-stage oocytes. (f) Percentage of SYCP3⁺ germ cells at the dictyate stage among total SYCP3⁺ cells at E18.5. Data are shown as mean +/− SD. \**P*<0.05, \*\**P*<0.01, \*\*\**P*<0.001 (two-tailed Student’s *t*-tests). Scale bars, 100 μm (overview) and 50 μm (higher magnification/insets).

At E14.5, analysis of germ cell development revealed a significant increase in SYCP3^+^ meiotic germ cells in anti-CSF1R-treated ovaries, while the number of TRA98^+^ total germ cells was grossly unaffected (Fig. 6c), indicating precocious meiotic entry without loss of germ cells. qRT-PCR analysis confirmed upregulation of meiosis-associated genes, including *Stra8, Sycp1, Sycp3, Syce1* and *Smc1b*, while the mRNA expression of *Pou5f1*and *Kit* (primordial germ cell and oogonia markers), as well as *Ddx4* (a general germ cell/oogonia marker), genes had no significant change (Fig. 6d). These results suggest that macrophage depletion accelerates germ cell meiotic differentiation, perhaps by altering the ovarian somatic environment during a sensitive developmental window.

By E18.5, we found that F4/80^+^ macrophages were restored in anti-CSF1R-treated ovaries, indicating a recovery of the macrophage population (Fig. 6e). However, the number of SYCP3^+^ germ cells in the dictyate stage remained significantly elevated (∼2-fold) compared to controls (Fig. 6f), demonstrating that the early depletion of macrophages caused a lasting advancement of meiotic progression. This suggests that transient loss of macrophages during a critical period is sufficient to induce long-term changes in germ cell behavior.

### Sustained macrophage depletion accelerates fetal ovarian germ cell meiotic progression

To specifically assess the long-term role of macrophages in fetal ovarian germ cell development, we utilized a *Csf1r*-Cre; *Csf1r^flox/flox^* knockout (*Csf1r*-KO) model, in which macrophages are ablated through targeted deletion of *Csf1r* in *Csf1r*-expressing cells (51). Compared to transient antibody-mediated CSF1R blockade, this genetic approach enables sustained macrophage depletion, allowing more precise study of macrophage function during ovarian development.

At E14.5, immunostaining confirmed a near-complete loss of F4/80^+^ macrophages in *Csf1r*-KO ovaries, while CD45^+^F4/80^−^ monocytes remained present, indicating specific depletion of macrophages (Fig. 7a). The total number of TRA98^+^ germ cells was not significantly changed between control (*Csf1r*-Cre; *Csf1r^flox/+^*) and *Csf1r*-KO ovaries (Fig. 7b and 7d), but the number of STRA8^+^ cells, as well as the percentage of STRA8^+^ germ cells, was significantly increased in *Csf1r*-KO ovaries (Fig. 7b, 7e, and 7f), indicating precocious meiotic initiation. Further staging of germ cells using SYCP3 and DDX4 showed a shift from the SYCP3^−^ (pre-meiotic) population to leptotene and zygotene stages in *Csf1r-*KO ovaries (Fig. 7c and 7g), indicating earlier meiotic entry following macrophage loss.

**Fig. 7.**
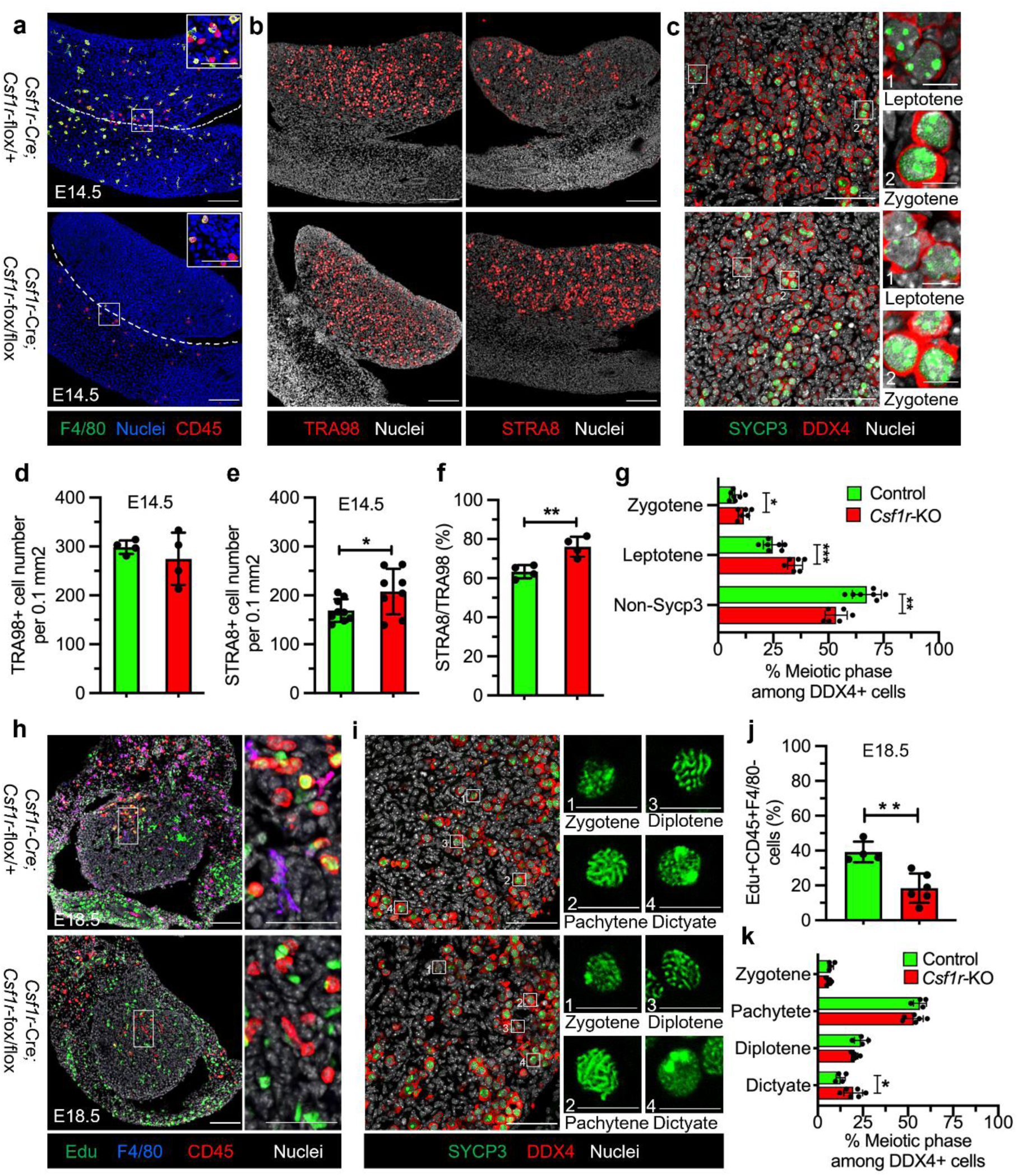
Genetic ablation of CSF1R⁺ macrophages leads to advanced fetal germ cell meiotic progression. (a) Representative E14.5 ovarian sections from control (*Csf1r*-Cre; *Csf1r^flox/+^*) (*n*=3) and *Csf1r*-KO (*Csf1r*-Cre; *Csf1r^flox/flox^*) (*n*=3) embryos stained for F4/80 and CD45, showing loss of macrophages but persistence of CD45⁺F4/80⁻ monocyte-like cells in *Csf1r*-KO ovaries. Insets show higher-magnification views of boxed regions; dashed lines mark the gonad-mesonephros boundary. (b) Representative E14.5 ovarian sections stained for TRA98 and STRA8 showing unchanged total germ cell distribution but increased STRA8⁺ cells in *Csf1r*-KO ovaries compared with control ovaries. For TRA98 staining, *n*=4 control and *n*=4 *Csf1r*-KO independent gonads; for STRA8 staining, *n*=9 control and *n*=8 *Csf1r*-KO independent gonads. (c) Higher-magnification images of E14.5 control (*n*=7) and *Csf1r*-KO (*n*=6) ovaries stained for SYCP3 and DDX4 to illustrate meiotic staging (non-SYCP3, leptotene, zygotene). (d-f) Quantification of TRA98⁺ germ cell number per 0.1 mm² area (d), STRA8⁺ germ cell number per 0.1 mm² (e), and percentage STRA8⁺ cells among TRA98⁺ cells (f) at E14.5. (g) Percentage of meiotic substages (non-SYCP3, leptotene, zygotene) among DDX4⁺ germ cells at E14.5. (h) Representative E18.5 sections stained for EdU, F4/80, and CD45, showing absence of F4/80⁺ macrophages and reduced EdU incorporation in CD45⁺F4/80⁻ cells in *Csf1r*-KO ovaries (*n*=6) compared with control ovaries (*n*=4). (i) E18.5 control (*n*=4) and *Csf1r*-KO (*n*=6) ovarian sections stained for SYCP3 and DDX4 to illustrate meiotic substages (zygotene, pachytene, diplotene, dictyate). (j) Percentage of EdU⁺ cells among CD45⁺F4/80⁻ monocyte-like cells at E18.5 in control and *Csf1r*-KO ovaries. (k) Percentage of meiotic substages among DDX4⁺ germ cells at E18.5 in control and *Csf1r*-KO ovaries. Data are shown as mean +/− SD. \**P*<0.05, \*\**P*<0.01, \*\*\**P*<0.001 (two-tailed Student’s *t*-tests). Scale bars, 100 μm (overview) and 50 μm (higher magnification/insets), except in c and i, where scale bars are 50 μm (overview) and 10 μm (higher magnification/insets).

At E18.5, *Csf1r*-KO ovaries showed a continued absence of F4/80^+^ macrophages, while CD45^+^ immune cells were still present (Fig. 7h). The proportion of CD45^+^F4/80^−^EdU^+^ monocytes was significantly reduced in *Csf1r*-KO ovaries compared to controls, indicating impaired monocyte proliferation or recruitment in the absence of CSF1R signaling (Fig. 7j). Concurrently, analysis of germ cell meiotic progression using SYCP3 and DDX4 co-staining showed that, although germ cells in both groups had entered late meiotic stages, a higher proportion of *Csf1r*-KO germ cells had reached the dictyate stage by E18,5, while control germ cells were more frequently in pachytene and diplotene stages (Fig. 7i and 7k). These results suggest that macrophage loss not only alters immune cell dynamics but also accelerates germ cell progression through meiotic prophase.

### Fetal macrophage loss disrupts perinatal ovarian germ cell clearance

Since *Csf1r-*KO mice die shortly after birth, limiting their utility for studying postnatal development, we applied an alternative approach to achieve sustained macrophage depletion. To further investigate the requirement for fetal ovarian macrophages—including those derived from the yolk sac and fetal HSCs—on postnatal ovarian development, we employed a long-term CSF1R blockade strategy by injecting pregnant mice with anti-CSF1R antibody at E6.5 and E14.5, which will ablate both yolk-sac-derived and HSC-derived macrophages. At E18.5, ovaries from antibody-treated embryos showed an almost complete loss of CD45⁺IBA1⁺ macrophages compared to controls (Fig. 8a), indicating efficient depletion of fetal macrophages. As development progressed, these macrophages gradually reappeared at P3 and returned to levels comparable to controls by P10 (Fig. 8a and 8c), suggesting robust postnatal repopulation. Interestingly, despite macrophage loss, the number of CD45⁺IBA1⁻ monocytes remained relatively stable compared to controls (Fig. 8a and 8d), indicating that the preservation of monocyte populations may contribute to the restoration of macrophages after birth. These findings suggest that while CSF1R-dependent macrophages are highly sensitive to depletion during fetal development, the ovary can effectively restore its macrophage population after birth. Thus, this dual anti-CSF1R antibody injection model provides a useful tool for studying the specific role of macrophages during early postnatal ovarian development.

**Fig. 8.**
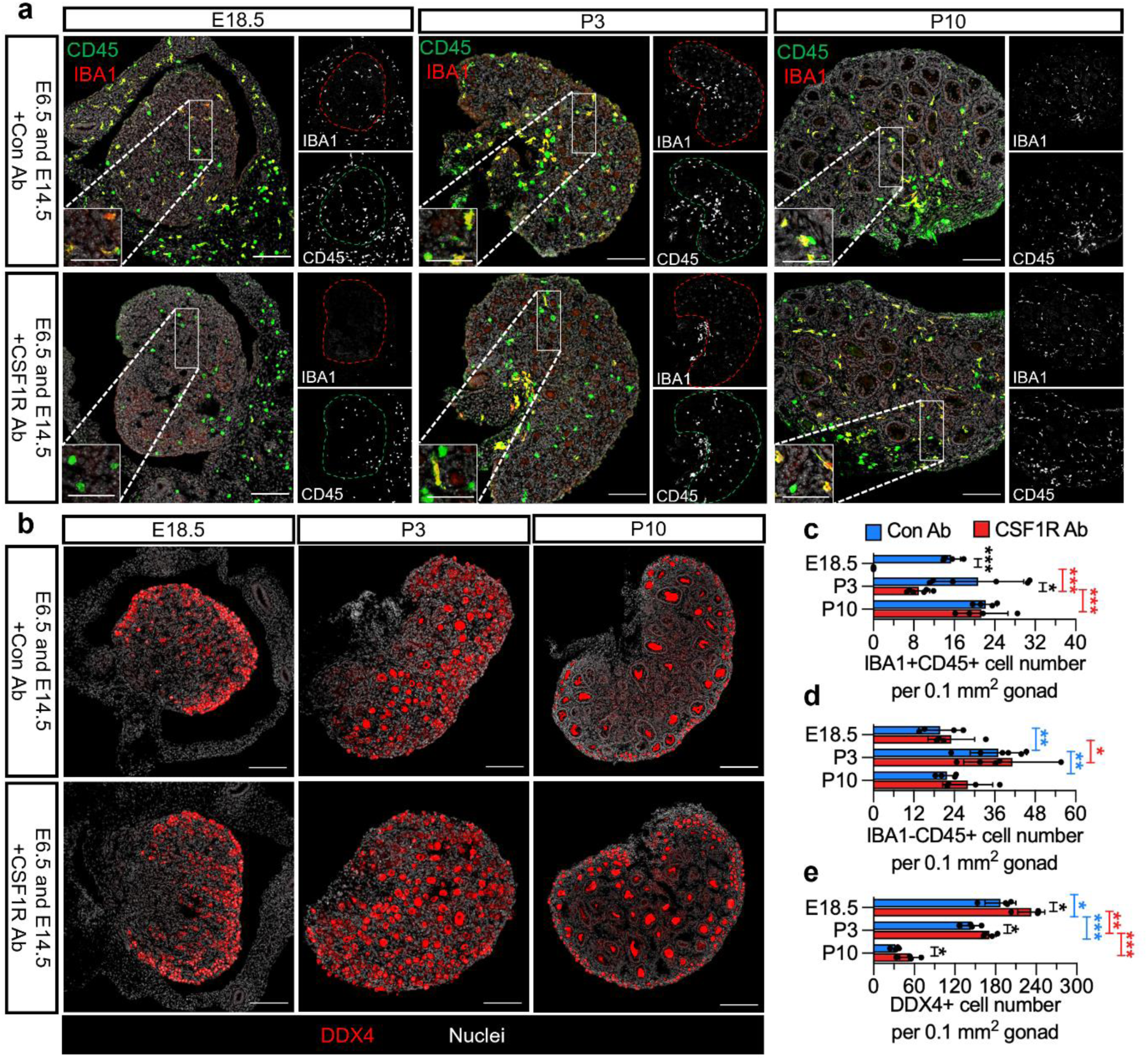
Fetal ovarian macrophage depletion impairs neonatal germ cell clearance while permitting postnatal macrophage repopulation. (a) Representative images of E18.5, P3, and P10 ovaries from C57BL/6J embryos exposed at E6.5 and E14.5 to either control rat IgG2a antibody or anti-CSF1R blocking antibody to deplete fetal ovarian macrophages. For E18.5, *n*=4 control and *n*=4 anti-CSF1R-treated independent gonads; for P3, *n*=6 control and *n*=6 anti-CSF1R-treated independent gonads; for P10, *n*=4 control and *n*=4 anti-CSF1R-treated independent gonads. Insets show higher-magnification views of boxed regions; red and green dashed outlines demarcate the ovary. (b) Representative sections stained for DDX4, showing developmental germ cell attrition at E18.5, P3, and P10 in control and anti-CSF1R-treated ovaries. For each time point, *n*=4 control and *n*=4 anti-CSF1R-treated independent gonads. (c) Quantification of CD45⁺IBA1^+^ macrophages per 0.1 mm² ovarian area at E18.5, P3, and P10. (d) Quantification of CD45⁺IBA1^−^ monocyte-like cells per 0.1 mm² ovarian area at the indicated stages. (e) Quantification of DDX4⁺ germ cell number per 0.1 mm² ovarian area at E18.5, P3, and P10, showing reduced physiological germ cell loss after macrophage depletion. Data are shown as mean +/− SD. \**P*<0.05, \*\**P*<0.01, \*\*\**P*<0.001 (two-tailed Student’s *t*-tests). Scale bars, 100 μm (overview) and 50 μm (higher magnification/insets).

In parallel, DDX4 staining showed that while the number of DDX4⁺ germ cells normally decrease from E18.5 to P10 as part of physiological germ cell attrition, this reduction was significantly attenuated in the anti-CSF1R-treated group. As a result, anti-CSF1R-treated ovaries contained more germ cells than controls at each stage examined (Fig. 8b and 8e). This suggests that macrophage depletion impairs the normal elimination of germ cells, possibly by disrupting the clearance of apoptotic cells or altering local inflammatory and paracrine signals that regulate germ cell selection. These findings highlight a role for CSF1R-dependent macrophages in facilitating developmental germ cell loss, which may be important for ensuring the quality and appropriate number of oocytes during ovarian maturation.

## Discussion

Our study delineates a developmental and functional framework for ovarian macrophages that integrates ontogeny, phenotypic heterogeneity, and germ cell regulation. We show that fetal ovarian macrophages arise from both yolk-sac-derived EMPs and fetal HSC-derived progenitors; that these embryonic cells persist postnatally; and that they are progressively complemented and replaced by CCR2-dependent blood monocytes, giving rise to three major MHCII/CSF1R-defined resident subsets. In parallel, CSF1R-dependent fetal macrophages restrain meiotic entry and promote physiological postnatal germ cell attrition, revealing a previously unappreciated developmental checkpoint controlled by phagocytes (52–54).

Previous work established that ovarian macrophages are abundant, broadly distributed, and essential for follicle development, ovulation, corpus luteum (CL) function, and tissue remodeling (4, 35, 55). Mass cytometry and fate mapping by Jokela et al. showed that fetal yolk-sac- and fetal-liver-derived macrophages coexist, persist after birth, and can convert from MHCII^−^ to MHCII^+^ states, but concluded that bone-marrow-derived monocytes contribute little to steady-state adult pools (39). Our data refine and extend this model by using temporally resolved *Csf1r*- and *Kit*-based lineage tracing, combined with *Cx3cr1*- and *Ccr2*-driven tools, to show a multi-wave contribution of EMPs, fetal HSCs, and CCR2^+^ monocytes to discrete MHCII/CSF1R subsets across fetal, postnatal, and adult stages. A recent study described F4/80^hi^CD11b^int^ and F4/80^int^CD11b^hi^ subsets with distinct CSF1R, MHCII, and CCR2 expression, as well as differing degrees of monocyte contribution and self-renewal (40). Additionally, single-cell analyses in mouse and human ovaries have highlighted transcriptional heterogeneity among ovarian myeloid cells, identifying multiple macrophage and monocyte clusters and age-associated shifts in ontogeny and polarization (56–58). Our work provides a developmental foundation for these transcriptomic states by mapping MHCII⁺CSF1R⁻, MHCII⁺CSF1R⁺, and MHCII⁻CSF1R⁺ populations onto distinct embryonic and monocyte-derived precursors and quantifying their proliferative dynamics from fetal life through adulthood. In contrast to earlier models that inferred ovarian ontogeny from surface markers and depletion surrogates (38, 40, 55), our lineage-tracing approach directly demonstrates that EMP-derived macrophages dominate in the fetal ovary; that fetal-HSC-derived progenitors become the principal source of adult macrophages; and that CCR2^+^ monocytes are specifically required for the maturation of MHCII⁺ subsets.

Comparisons with other organs underscore both shared and ovary-specific principles of macrophage biology. In multiple tissues, including heart, pancreas and gut, embryonic macrophages are progressively replaced by adult monocyte-derived cells, ontogenically distinct subsets that exhibit different turnover kinetics under steady-state conditions (59–61). Ovarian macrophages follow this general scheme, but our data and prior work indicate distinctive kinetics and subset relationships: fetal EMP-derived cells initially seed the ovary; fetal-liver/HSC-derived cells expand during late gestation; and postnatal CCR2^+^ monocytes preferentially generate MHCII⁺ populations that become dominant with age. This stands in contrast to the testis, where yolk-sac-derived macrophages contribute minimally to adult populations, and a narrow embryonic window of HSC-derived monocyte recruitment builds long-lived interstitial and peritubular macrophages that promote organogenesis, Leydig-cell steroidogenesis, and spermatogonial differentiation (20, 21, 62, 63). Together, these comparisons reveal that gonads share a reliance on fetal and monocyte-derived macrophages, but they deploy distinct developmental programs and functional specialization in the ovary versus testis.

Functionally, our work adds a germ-cell-centric dimension to the established roles of ovarian macrophages. Previous studies have shown that macrophages regulate folliculogenesis, ovulation, corpus luteum (CL) development, and vascular integrity, and that experimental depletion using clodronate liposomes, CD11b-DTR, or CSF1/CSF1R blockade impairs ovulation, destabilizes luteal vasculature, alters steroidogenesis, and compromises implantation (55, 64–67). Single-cell and imaging studies further indicate that ovarian macrophages and related phagocytes are enriched around regressing follicles and degenerating oocytes, and that they change their activation state with age and cycle stage, supporting a role in follicle atresia, tissue remodeling, and “inflammaging” (38, 56, 58). Beyond the ovary, testicular macrophages are proposed to be integral components of the spermatogonial niche that supports germ cell differentiation (20), while epididymal immune cells, including dendritic cells and mononuclear phagocytes, contribute to the clearance of defective or apoptotic germ cells and maintenance of sperm quality (68, 69). These findings support a broader concept that reproductive tract phagocytes participate directly in germ cell quality control.

Several lines of evidence directly link innate immune signaling to germ-cell-intrinsic cell cycle control. In the testis, macrophage inflammatory protein-1α (MIP-1α/CCL3) acts locally to stimulate DNA synthesis in primitive spermatogonia and premeiotic spermatocytes while inhibiting more differentiated spermatogonia, indicating that a macrophage-derived chemokine can differentially tune mitotic and meiotic progression of germ cells (70). In the mammalian ovary, CSF1 produced by granulosa and immune cells participates in LH-triggered meiotic resumption: CSF1 signaling through its receptor (CSF1R) modulates expression of the NPPC/NPR2 axis in preovulatory follicles, lowers NPR2 levels, and promotes oocyte germinal-vesicle breakdown, thereby acting as an intermediary between gonadotropin signaling and oocyte meiosis (71, 72). Complementing these mammalian data, zebrafish studies have established a functional link between macrophage activation and germ cell loss: in *Bmp*15 mutants, definitive CSF1R-dependent macrophages are required for pathological oocyte depletion, ovarian failure, and female-to-male sex reversal, and genetic ablation of macrophages or Csf1rb ligands (Il34/Csf1a) delays or prevents oocyte loss and masculinization (34). Together, these findings across species support the concept that innate immune cells do not simply clear debris but actively direct germ cell fate, meiotic timing, and gonadal development. Against this background, our data show that CSF1R-dependent fetal ovarian macrophages restrain the timing of meiotic initiation and are required for efficient postnatal germ cell clearance. By acting before and during the wave of physiological oocyte attrition, these embryonically imprinted macrophages function as gatekeepers of the oocyte pool, enforcing a quality-control checkpoint that may operate in parallel with somatic pathways such as the LH-CSF1-NPPC/NPR2 signaling cascade described above.

Our findings also indicate that ontogenically distinct macrophage subsets execute non-redundant functions within this germ cell regulatory network. EMP- and fetal HSC-derived macrophages, enriched among early MHCII⁻CSF1R⁺ cells, colonize the fetal ovary when germ cells enter meiosis and are anatomically positioned along vessels and stromal interfaces, akin to embryonic testicular macrophages that coordinate cord formation and vascular remodeling to permit normal spermatogonial development (20, 73). In contrast, monocyte-derived MHCII⁺ populations, which expand postnatally in a *Ccr2*-dependent manner, may be better suited to long-term tissue remodeling and age-associated inflammation. This is consistent with scRNA-seq and single-cell atlas studies showing that ovarian myeloid compartments diversify with age, with accumulation of pyroptotic and profibrotic macrophage states, and remodeling of stromal and granulosa niches during ovarian aging (57, 58, 74). Together, these data argue that the timing and balance of embryonic versus monocyte-derived macrophages are critical variables when interpreting immune contributions to ovarian physiology and pathology, and they provide a mechanistic context in which a fetal macrophage-dependent meiotic checkpoint can be understood.

## Methods

### Mice

All animals were maintained under specific pathogen-free conditions at the animal facility of Cincinnati Children’s Hospital Medical Center (CCHMC) under a 12 h light/12 h dark cycle at 22 °C and 40-60% humidity, with ad libitum access to food and water. All procedures conformed to institutional and National Institutes of Health guidelines and were approved by the CCHMC Institutional Animal Care and Use Committee (IACUC protocols IACUC2018-0027, IACUC2021-0016, and IACUC2024-0039). The following strains were used in this study: *Csf1r*-creER [Tg(Csf1r-Mer-iCre-Mer)1Jwp/J; JAX #019098] (48), *Cx3cr1*-creER [Cx3cr1^tm2.1(creERT2)Jung^/J; JAX #020940] (46), *Csf1r*-Cre [Tg(Csf1r-icre)1Jwp/J; JAX #021024] (75), *Ccr2*-GFP [Ccr2^tm1.1Cln^/J; JAX #027619] (76), *Rosa*-Tomato [Gt(ROSA)26Sor^tm14(CAG-tdTomato)Hze^/J; JAX #007914] (77), *Csf1r*-flox [Csf1r^tm1.2Jwp^/J; JAX# 021212] (78) and wild-type C57BL/6J (B6) (JAX #000664) mice. *Kit*-creER mice [Kit^tm2.1(cre/Esr1*)Jmol^/J] (79) were generated by J. Molkentin at Cincinnati Children’s Hospital Medical Center and are commercially available from The Jackson Laboratory (JAX stock #032052). All strains were maintained on a C57BL/6J background unless otherwise noted. Both male and female breeders were 8-16 weeks of age. Embryonic stages were defined by timed matings, with noon on the day of vaginal plug detection designated as embryonic day (E) 0.5.

### Tamoxifen-induced lineage tracing

To trace the developmental origins of ovarian macrophages, we used *Csf1r*-creER; *Rosa*-Tomato, *Kit*-creER; *Rosa*-Tomato, and *Cx3cr1*-creER; *Rosa*-Tomato lineage-tracing models. Pregnant females were injected intraperitoneally at defined stages with 4-hydroxytamoxifen (4-OHT, Sigma-Aldrich #H6278) dissolved in corn oil and supplemented with progesterone subcutaneously (Sigma-Aldrich #P0130) to limit pregnancy loss. For embryonic labeling, *Csf1r*-creER; *Rosa*-Tomato and *Kit*-creER; *Rosa*-Tomato pregnancies received 75 µg/g body weight 4-OHT plus 37.5 µg/g body weight progesterone at E8.5, E10.5, or E12.5. For *Cx3cr1*-creER; *Rosa*-Tomato experiments, two regimens were used: to label embryonic *Cx3cr1*^+^ macrophages, pregnant females were injected with 75 µg/g 4-OHT and 37.5 µg/g progesterone at E12.5; to label early postnatal monocytes and macrophages, *Cx3cr1*-creER; *Rosa*-Tomato pups received 50 µg tamoxifen (TAM, Sigma-Aldrich #T5648) intraperitoneally on P4 and P5.

### Antibody-mediated depletion of CSF1R^+^ macrophages

To transiently deplete yolk-sac-derived and fetal CSF1R^+^ macrophages, pregnant C57BL/6J females were injected intraperitoneally with 3 mg anti-CSF1R monoclonal antibody (mAb; clone AFS98, Bio X Cell #BP0213) or rat IgG2a isotype control (Bio X Cell #BP0089). For early depletion studies as previously described (9, 21), anti-CSF1R or control IgG antibody was administered at E6.5 and ovaries were collected at E13.5, E14.5, and E18.5. This regimen targets yolk-sac-derived macrophages and their progeny. To extend depletion into late fetal stages and evaluate consequences for neonatal ovarian development, a second cohort of pregnant females received anti-CSF1R or control IgG injections at both E6.5 and E14.5. Ovaries from these litters were collected at E18.5, P3, and P10 to assess macrophage repopulation and germ cell attrition.

### EdU incorporation in vivo

Cell proliferation was examined by 5-ethynyl-2′-deoxyuridine (EdU) incorporation. Pregnant females or postnatal pups received an intraperitoneal injection of EdU (Invitrogen Click-iT EdU Alexa Fluor 488 kit, #C10337) at a dose of 25 μg/g of body weight diluted in sterile PBS. For embryonic analyses (E14.5 and E18.5), pregnant females were injected 4 h before embryo collection. For postnatal stages, pups were injected 2 h prior to euthanasia. Ovaries were processed for cryosectioning, and EdU detection was performed using the Click-iT reaction according to the manufacturer’s instructions, followed by immunostaining as described below. Proliferating macrophage and monocyte-like populations were quantified as EdU⁺ cells within defined immunophenotypic gates.

### Immunofluorescence

For whole-mount immunofluorescence, E12.5 and E13.5 ovaries with attached mesonephroi were dissected in PBS and fixed overnight at 4°C in 4% paraformaldehyde (PFA) containing 0.1% Triton X-100, as previously reported (80). After fixation, tissues were washed in PBS with 0.1% Triton X-100 (PBTx), blocked in PBTx supplemented with 10% FBS and 3% bovine serum albumin (BSA) for 1-2 h at room temperature, and incubated with primary antibodies diluted in blocking solution overnight at 4°C.

For later embryonic and postnatal stages (E14.5, E16.5, E18.5, P3, P10, P14, P30, P60, P90), ovaries were fixed overnight in 4% PFA with 0.1% Triton X-100 at 4°C, cryoprotected in a sucrose gradient series (10%, 15%, 20% sucrose in PBS), embedded in OCT compound (Fisher Healthcare #4585), and frozen at −80 °C. Cryosections (typically 10-14µm) were collected on glass slides, rehydrated in PBTx, blocked in PBTx with 10% FBS and 3% BSA, and incubated with primary antibodies overnight at 4°C.

Primary antibodies included markers for macrophages and monocytes (F4/80, IBA1, CD45, MHCII, CSF1R), endothelial cells (PECAM1/CD31), and germ cells and meiotic stages (TRA98, DDX4, STRA8, SYCP3). Detailed antibody information is provided in Supplementary Table 1. After several washes in PBTx, samples were incubated with species-appropriate Alexa Fluor-conjugated secondary antibodies (Alexa Fluor 488, 555, or 647; Thermo Fisher; 1:500) and Hoechst 33342 nuclear dye (1 mg/ml; Thermo Fisher #H1399) for 1 h (sections) or 2-3 h (whole mounts) at room temperature. All nuclear stains were performed using Hoechst 33342 and are labeled as “Nuclei” within figure panels. Images were acquired on a Nikon Eclipse TE2000 microscope equipped with structured illumination (OptiGrid/Volocity; PerkinElmer) or on a Nikon A1 inverted confocal microscope (Nikon). Acquisition settings were kept constant within each experiment.

### Quantitative real-time PCR (qRT-PCR)

Total RNA was isolated from pooled ovaries of defined stages and treatment groups using TRIzol reagent (Invitrogen #15596018) according to the manufacturer’s protocol as previously described (21). Residual genomic DNA was removed with DNase I (Amplification Grade, Thermo Fisher #18068015). cDNA was synthesized from 500 ng RNA using an iScript cDNA synthesis kit (Bio-Rad #1708841). qRT-PCR was performed using Fast SYBR Green Master Mix (Applied Biosystems #4385616) on a StepOnePlus real-time PCR system (Applied Biosystems). Gene expression was normalized to *Gapdh* and relative fold changes were calculated using the 2^−ΔΔCt^ method. Primer sequences for macrophage-associated genes, endothelial marker, and germ cell/meiotic regulators are listed in Supplementary Table 2. At least three independent biological replicates (each consisting of pooled ovaries from ≥2 embryos or pups) were analyzed per condition.

### Quantification of ovarian cell populations

Fetal and adult ovarian macrophages/immune cells, lineage-traced Tomato⁺ cells, and germ cells were quantified on serial ovarian sections or whole-mount preparations at the indicated developmental stages. For each ovary, all sections containing ovarian tissue were collected at a uniform thickness, and every second or third section was selected for analysis depending on ovary size. Cells of interest were identified based on fluorescence and morphology and were manually counted using the Cell Counter plugin in ImageJ/Fiji (NIH). Total cell number per ovary was estimated by multiplying the counted cell number by the section sampling factor. Ovarian area was measured using the polygon selection tool in ImageJ to allow normalization where indicated.

To evaluate meiotic entry and progression, germ cells were classified into pre-meiotic and sequential meiotic substages (leptotene, zygotene, pachytene, diplotene/dictyate) according to nuclear morphology and chromosome axis organization visualized by immunostaining, based on established criteria (81). For each ovary, at least three separate sections were analyzed. All counting and staging were performed in a genotype- and treatment-blinded manner.

### Statistics and reproducibility

For all experiments, *n* refers to independent biological replicates (individual embryos or pups, or pooled ovaries from a single litter when indicated). For immunofluorescence analyses, at least three sections from each of *n*≥3 independent animals were examined per group. For qRT-PCR, data represent 2-3 independent experiments, each with *n*≥3 biological replicates.

Graphing and statistical analyses were performed using GraphPad Prism (version 8 or later). Data are presented as mean ± SD unless otherwise noted. Comparisons between two groups were made using unpaired two-tailed Student’s *t*-tests. Exact tests, *n* values, and *P*-values are reported in the figure legends.

## Author Contributions

XG and TD designed research; XG, SM, S-YL, and TD performed research; XG and TD analyzed data; XG, SM, S-YL, and TD edited the paper; and XG and TD wrote the paper.

## Acknowledgements

We thank Dr. Jeffery Molkentin for *Kit*-creER mice and Dr. Dagmar Wilhelm for anti-FOXL2 antibody. This work was supported by National Institutes of Health (R35GM119458 to TD) and Cincinnati Children’s Hospital Research Foundation (Research Innovation and Pilot Funding to TD). We also acknowledge BioRender (BioRender.com) for the use of images.

## Supplementary Tables

**Supplementary Table 1.** Primary antibodies used for immunofluorescence (IF)

| Primary Antibody (IF) | Dilution | Source/Reference |
| --- | --- | --- |
| Rat anti-CD11b | 1:250 | BD Pharmingen #557395 |
| Goat anti-CD45 | 1:500 | R&D #AF114 |
| Rat anti-CD45 | 1:300 | BioLegend #103101 |
| Rabbit anti-CSF1R | 1:500 | CST #43390S |
| Rabbit anti-DDX4 | 1:1,000 | Abcam #ab13840 |
| Rat anti-F4/80 | 1:2,000 | AbD Serotec #MCA497RT |
| Rabbit anti-FOXL2 | 1:500 | D. Wilhelm (1) |
| Chicken anti-GFP | 1:1,000 | Aves #GFP-1020 |
| Rabbit anti-IBA1 | 1:1,000 | Wako #019-19741 |
| Rat anti-MHCII | 1:500 | eBioscience #14-5321-81 |
| Goat anti-PECAM1 | 1:250 | R&D #AF3628 |
| Rat anti-PECAM1 | 1:250 | BD Pharmingen #553370 |
| Rabbit anti-STRA8 | 1:2000 | Abcam #ab49602 |
| Mouse anti-SYCP3 | 1:500 | Abcam # ab97672 |
| Rat anti-TRA98 | 1:1,000 | Abcam #ab82527 |

**Supplementary Table 2.**
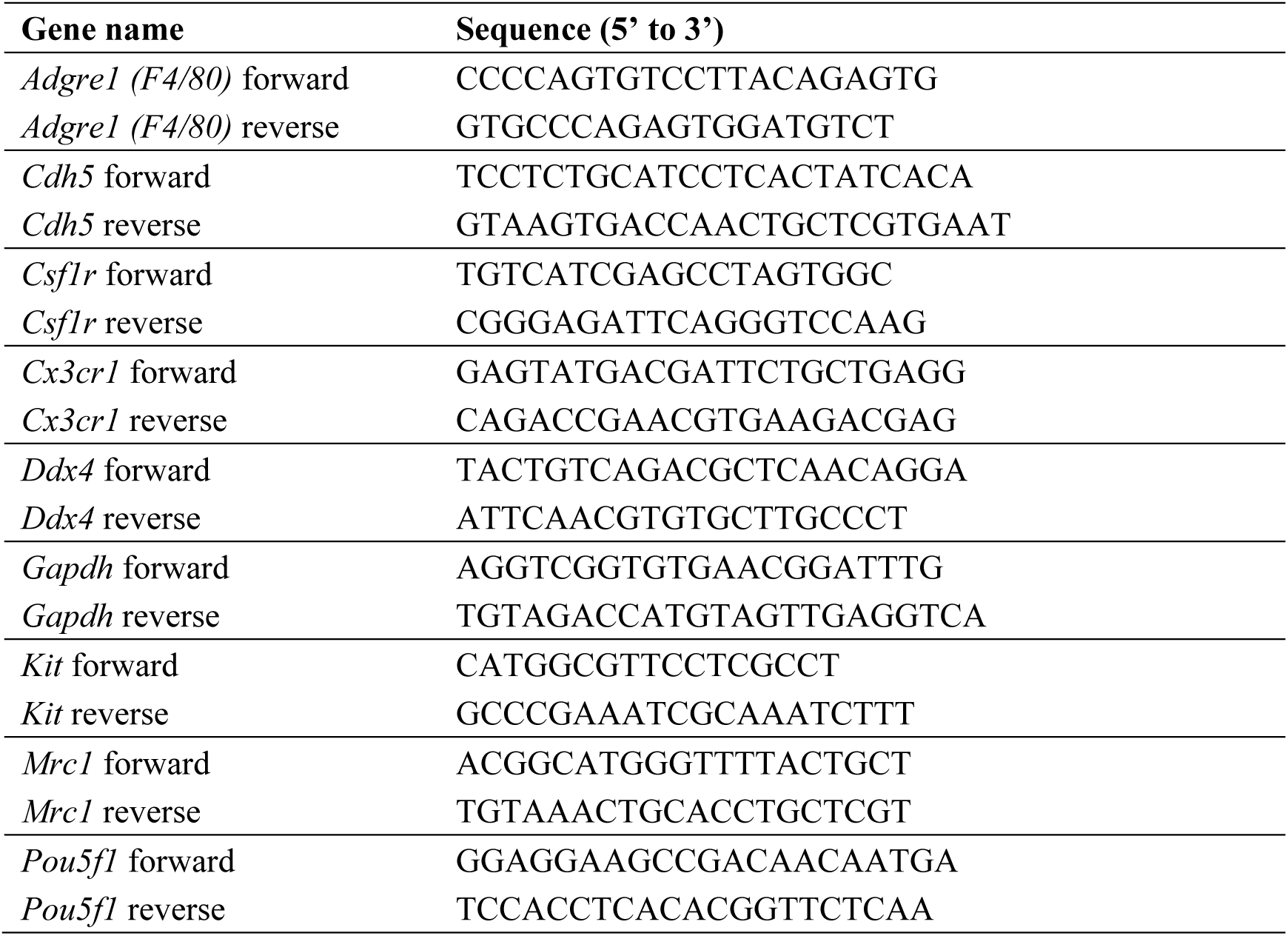
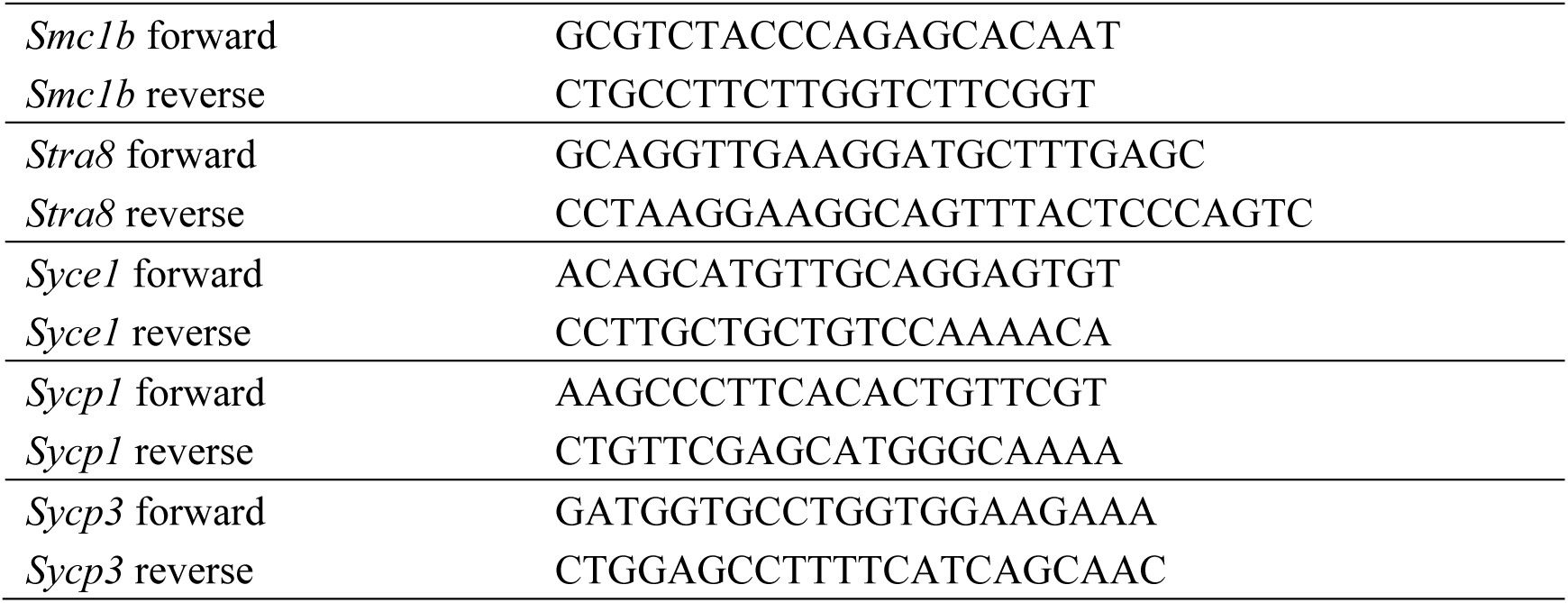
Sequences of primers used for qRT-PCR analyses.

